# Codon-Dependent Translational Accuracy Controls Protein Quality in *Escherichia coli* but not in *Saccharomyces cerevisiae*

**DOI:** 10.1101/200006

**Authors:** Lyne Jossé, Connor D. D. Sampson, Mick F. Tuite, Kevin Howland, Tobias von der Haar

**Affiliations:** Kent Fungal Group, School of Biosciences, University of Kent, Canterbury, CT2 7NJ, UK

## Abstract

In order to generate a functional proteome, gene expression pathways must assemble proteins accurately according to the rules of the genetic code. General gene expression accuracy is known to be high, but errors nevertheless occur with measurable frequencies. Here we develop a mass-spectrometry (MS) based assay for the detection of a particular type of gene expression error, amino acid misincorporation. This assay allows assessing a much broader range of misincorporation events compared to current, very sensitive but also very specific enzyme reporter assays. Our assay uncovers a remarkably rich pool of error products for a model protein expressed in *E. coli*, which depend quantitatively on codon usage in the expression construct. This codon usage dependence can be explained in part as a function of the composition of the tRNA pool in this organism. We further show that codon-dependent differences in error levels correlate with measurable changes in specific protein activity. In contrast to *E. coli*, error levels are lower, and appear not to be codon usage dependent, when the same model protein is expressed in *S. cerevisiae*.

## Introduction

The accurate synthesis of proteins according to information contained in the DNA code is essential for all biological processes. However, protein synthesis accuracy is not infinite, and various types of error, including random misincorporation of amino acids, occur with low but measurable frequencies ^1,2^. Such errors can have various outcomes depending on the affected site and the nature of the misincorporated amino acid, but typically they tend to reduce the functionality of the affected proteins^3^. Cells can adapt to increased errors, but this generally comes at the price of increased energy expenditure^4,5^. Error levels above the normal physiological range interfere with biological processes and produce physiological deficiencies like mitochondrial dysfunction ^6^, accelerated cellular ageing ^7^ and disease conditions^8^.

Amino acid misincorporation can occur due to a number of causes, including transcriptional errors^9^, errors during tRNA aminoacylation^10^, or during tRNA selection by the ribosome^11^ (Figure 1A). In all these cases, information processing molecules (RNA polymerases, tRNA synthetases or ribosomes) must distinguish between competing appropriate and inappropriate molecular entities. The accuracy of these processes depends on both the physical similarity of the competing species, and on their relative levels. Especially for tRNA selection, this principle has been well established through *in vivo* experimentation ^11^.

**Figure 1.**
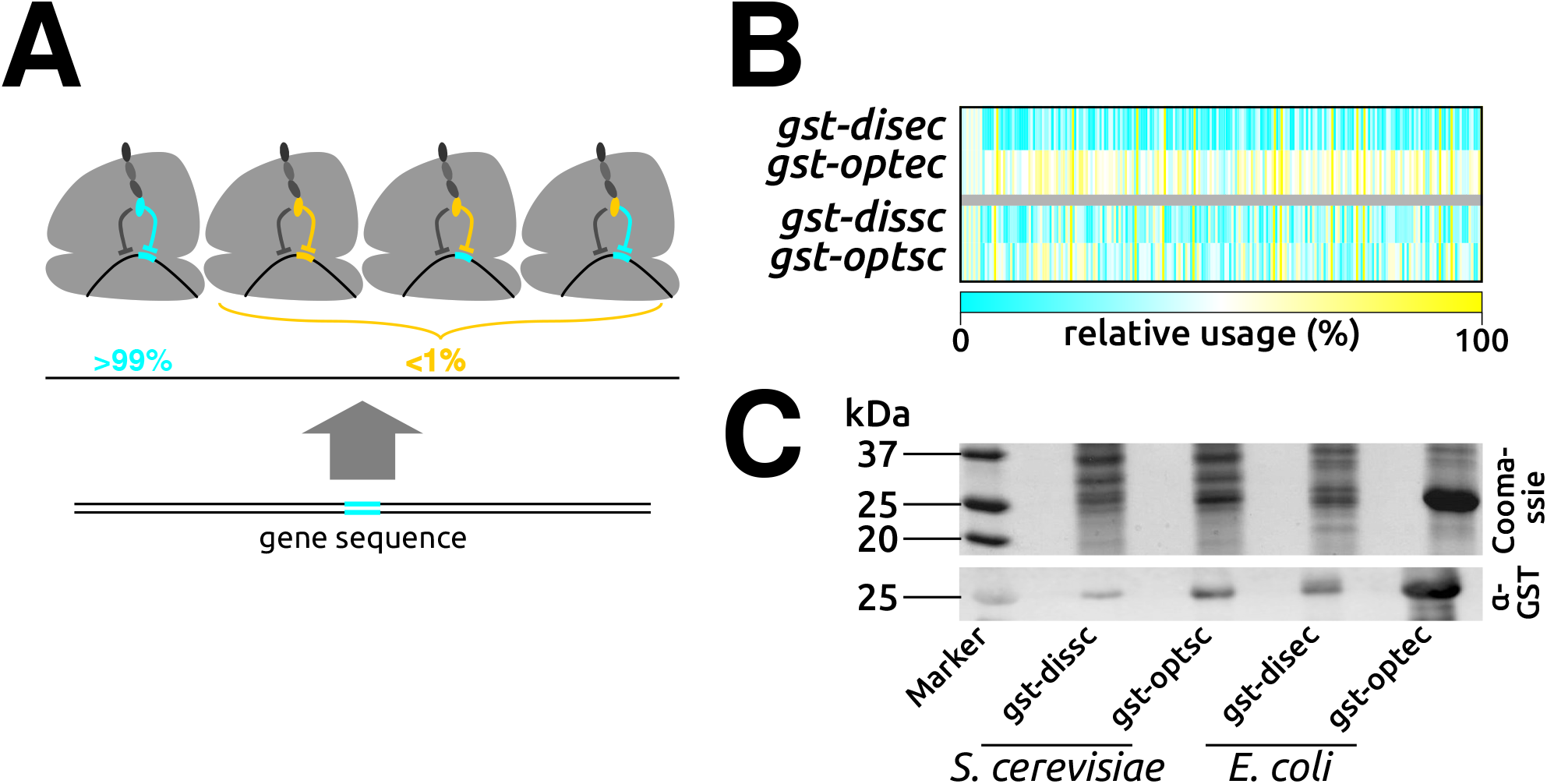
Gene expression errors. **A**, accurate decoding requires the matching of amino acids to their correct tRNA, and matching of tRNAs to their correct codon. Errors can occur if transcription alters the codon relative to the DNA sequence, if the ribosome pairs the incorrect tRNA with the codon, or if tRNA synthetases connect the incorrect amino acid to a tRNA. **B**, illustration of codon usage differences in the GST expression constructs used in this study. Colour identifies the relative usage of individual codons from rarely used (blue) to frequently used (yellow). **C**, expression levels of the different GST alleles used in this study.

Gene expression accuracy is achieved through evolutionary optimisation, which ensures that errors occur as infrequently as possible while still allowing gene expression rates that are compatible with cellular requirements ^12^. Several lines of evidence suggest that physiological gene expression accuracy has evolved to a point where error levels are just low enough to avoid interfering with biological processes. Relevant observations include the apparent co-evolution of longevity and translational accuracy in mammals ^13^, and the evolutionary selection for less error-prone codons at structurally sensitive sites in proteins^14^. Moreover, several studies observed apparent regulation of amino acid misincorporation levels by cellular signalling pathways ^7,15,16^. This apparent close matching of physiological accuracy to cellular requirements raises the question in how far accuracy can be maintained under conditions where evolutionary optimisation is reduced, such as non-evolved recombinant protein expression systems, or in cells in which optimisation is lost because of diseases.

Published studies of amino acid misincorporation errors have relied on sensitive reporter systems for measuring such errors. In particular, the recovery of enzymatic activity in enzyme dead mutants of reporter constructs like chloramphenicol acetyl transferase^17^, β-galactosidase^18^ or luciferases^19,20^ has substantially advanced the field. However, these constructs only allow assessing a limited selection of the 20x19 theoretically possible amino acid misincorporation events, and consequently there is a need for novel assays for studying a broader range of errors. Mass spectrometric approaches are emerging as one promising avenue for developing such assays^4,21,22^. To date such assays have allowed detecting amino acid misincorporation under conditions of reduced accuracy, but to our knowledge they have not yet been used to detect physiological errors. Here we describe a mass spectrometric strategy which we apply to reporter proteins produced from different coding sequences in *Echerichia coli* and *Saccharomyces cerevisiae*. Our assay reveals a rich repertoire of apparent amino acid substitution events, which in *E. coli* but not in yeast are quantitatively dependent on the coding sequence used to express the protein. We also show that physiological, sequence dependent translational errors in *E. coli* correlate with measurable differences in protein activity.

## Results

### An experimental system for studying amino acid misincorporation errors *in vivo*

We initially constructed a series of expression vectors for a histidine-tagged glutathione-S-transferase (GST). GST is easily and highly expressed in different organisms, and the tag enables crude purification without relying on the endogenous affinity of the enzyme for glutathione, which could skew the composition of mixed native and mis-translated GST populations.

We constructed genes encoding identical 8xHis-GST proteins by distinct DNA sequences, that differed in the codons used to encode the GST part of the protein. We designed two sequences for expression in *E. coli*, consisting of the most and least preferred codons based on published codon frequencies in this organism^23^. Two further sequences were designed for expression in yeast, consisting of the fastest and slowest decoded codons according to our published decoding models ^24^ (figure 1B). We refer to these sequences as *gst-optec* and *gst-disec* for *E. coli*, and *gst-optsc* and *gst-dissc* for *S. cerevisiae*^i^.

Although there is a wide-spread assumption in the literature that preferred codons should result in low error levels and non-preferred codons in high levels, Shah and Gilchrist ^25^ noted that this rule may not hold for organisms in which the abundance of cognate and near-cognate tRNAs is correlated. Thus, we did not necessarily assume that the two DNA sequences for each organism would generate proteins with differing bulk error levels, but we reasoned that we might observe qualitative differences in misincorporation patterns for the different sequences and hosts.

Upon expression in their respective host organisms, the different coding sequences yielded substantially differing protein levels (figure 1C), as was expected from existing studies on codon optimisation of recombinant constructs in these organisms ^24,26^. To analyse amino acid misincorporation in the different expression products by mass spectrometry, we prepared samples that were semi-purified via immobilised metal affinity chromatography (IMAC). These samples were then separated on SDS-PAGE gels, with the sample loading volume adjusted so that the bands representing GST were of roughly equal strength. The bands were excised, subjected to in-gel tryptic digest, and the resulting tryptic peptides analysed by mass spectrometry using a nano-LC-MS/MS setup. By utilising a data-independent acquisition (DIA) approach to the analyses, we were able to collect MS/MS data on low abundance peptides in the presence of more abundant peptides. The DIA approach utilised an additional gas-phase separation based on the collisional cross section of the ions (ion mobility separation [IMS]), which gave additional resolution of the complex samples. Coupled with alternating low and high energy MS scans, his allowed us to generate MS/MS spectra for high numbers of peptides eluting at any given ^27,28^.

To detect translation products resulting from amino acid misincorporation events, we produced a systematic list of all tryptic peptides that could result from single amino acid substitutions in the GST sequence. This list included substitutions from K and R to other amino acids and *vice versa*, which alter tryptic digest patterns. The list was used to search the MS/MS spectra generated as described above.

### Detection of peptides containing *bona fide* amino acid misincorporation errors

Our mass spectrometry work flow relied on sensitive detection methods, which carry the risk of false discovery of individual peptides. The Progenesis QIP software (Nonlinear Dynamics) used to evaluate mass spectra in our work includes calculation of false peptide detection rates in its calculation of the detection scores, and we restricted our subsequent analyses to peptides with a score > 5 where false discovery rates should be minimal. However, as we were using a non-standard experimental strategy, it is not clear in how far the false detection rate calculations are really accurate in our setting. We therefore sought to extensively validate the MS/MS data by asking in how far properties of the detected peptides made sense in the light of known or predicted biological properties of amino acid misincorporation peptides. In the first instance, we analysed pooled data generated with the different expression sequences and hosts and from multiple experiments.

We detected 9 of the 30 possible tryptic peptides derived from the wild-type GST sequence (“wild type peptides”). Additionally, we detected a large number of peptides corresponding to single amino acid substitutions in the GST sequence (“error peptides”). In all we detected 101 apparent substitution events at 62 distinct sites (figure 2A). Most of the detected error peptides had corresponding wt peptides that were also detected, although for seven error peptides this was not the case. Conversely, for four wt peptides, no corresponding error peptide was detected. For substitutions that introduced or removed lysine or arginine residues, we observed the expected corresponding changes in the tryptic cleavage patterns.

**Figure 2.**
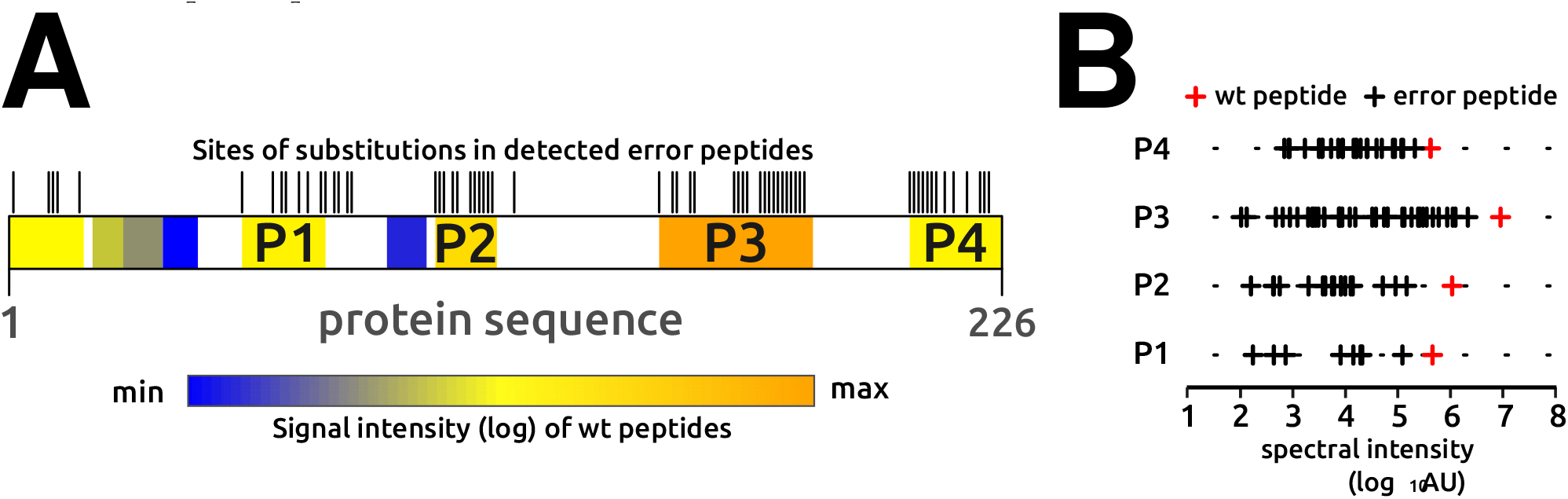
Mass spectrometric detection of wild-type and error peptides. **A** shows a representation of the primary sequence of GST, indicating in colour the location of peptides for which the expected wild-type sequence was detected in MS/MS experiments. The colour indicates the relative strength of the spectral signal for each peptide. Ticks above the boxes indicate sites of detected amino acid substitutions. **B**, signal intensities of the wild-type peptides (red) and their derived error-containing peptides (back). See main text for discussion.

The number of detected error peptides correlated strongly with the signal intensity observed for the corresponding wild-type peptide, and the typical spectral intensity of error peptides was orders of magnitude lower than the intensity of the corresponding wt peptides (figure 2B). These analyses are limited because our mass spectrometric experiments were not strictly quantitative, but the data are consistent with the expectation that errors are rare and error peptides make up small sub-populations of the total peptide mixture.

Analyses of the observed single substitutions by type reveal that, of the 20x19 possible substitutions, 76 (20%) were observed (figure 3A). 12 substitutions were observed at multiple independent sites, with five (A→N, A→S, D→M, L→H and P→E) observed at four independent sites each. Given the total number of substitutions we detect, observing five different substitutions at four independent sites each would occur by chance with a frequency of less than 1 in 10^9^. Thus, the observed substitutions are very likely linked to underlying biological or chemical mechanisms.

**Figure 3.**
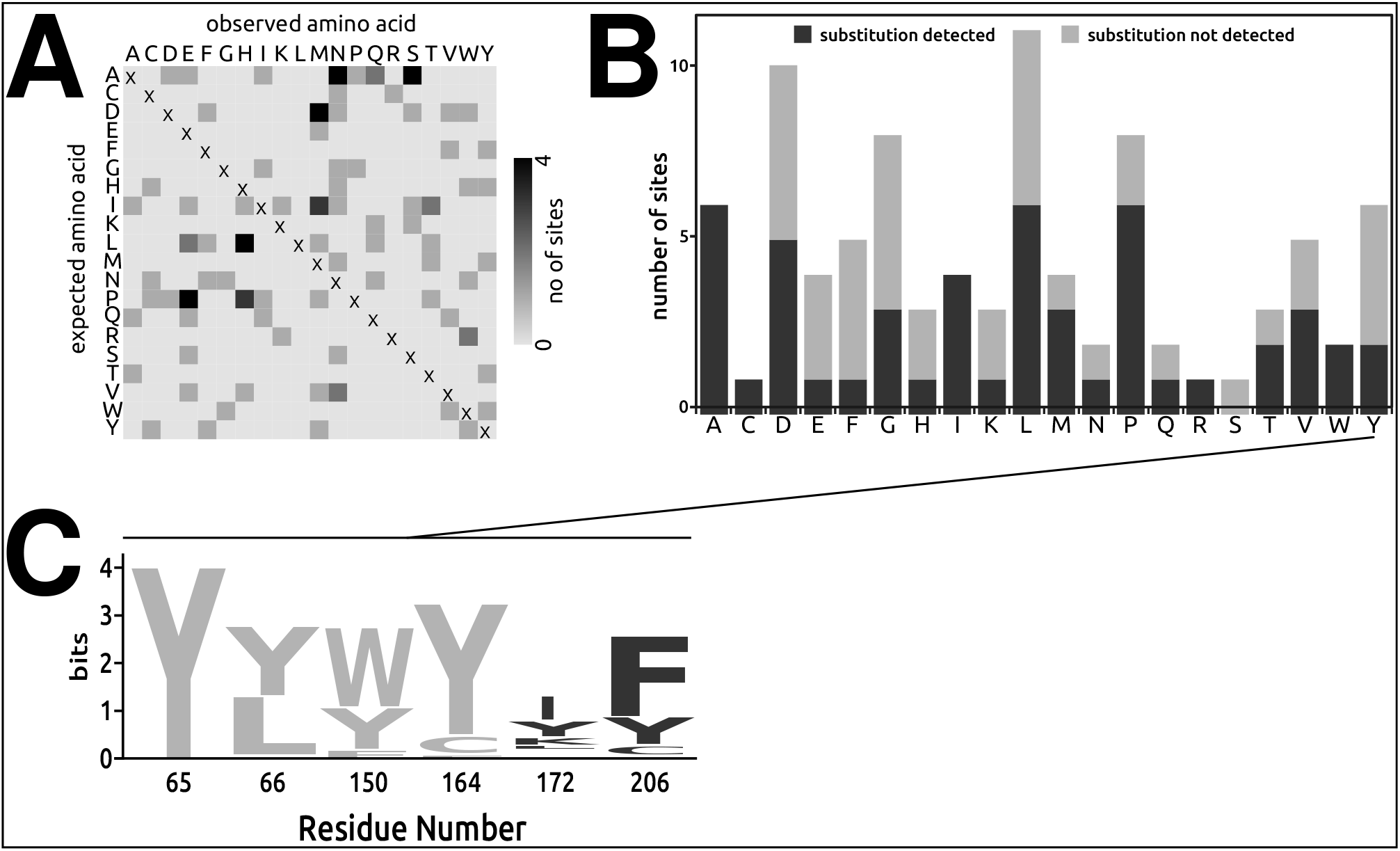
Detected substitution events tend to not affect protein stability. **A**, summary of detected substitutions. Individual substitutions were detected on up to four independent sites in the GST sequence. **B**, comparison of sites available in the detected wild type peptides (total bar height), and the number of sites for which substitutions were detected (dark grey portion of bars). **C**, sequence variability in the 50 most closely related homologues of GST in UNIREF-90, for the example of tyrosine codons. Detected substitutions are Y→F, Y→M, and Y→W, at site 172, and Y→C at site 206, indicating that substituted peptides are preferentially detected at sites where amino acid changes are well tolerated.

We also attempted to detect double error peptides, by generating an exhaustive list of double substitutions combining single substitutions observed in in the well detected peptide P3. The original ion search revealed 29 individual substitutions in this peptide, generating a potential for 334 detectable double substitutions (this is less than the mathematically possible 29x28 double substitutions because substitutions for different amino acids at the same site cannot be combined). We were only able to detect signals for 14 of these double substitutions, consistent with the notion that error probabilities are independent of each other, and peptides with multiple errors are therefore becoming exponentially less abundant and harder to detect.

The synthetic genes used in this study consist of one single codon type for each of the amino acids, meaning that if translational errors occur at one site, they should be equally likely at all other sites. For alanine, isoleucine and tryptophan we do indeed observe substitutions at all available sites within the detected peptides (figure 3B) (arginine, cysteine and serine occur only at a single site within the covered peptides). However, for the remaining 14 amino acids we observed substitutions at some sites but not others. We reasoned that this could be explained if translational errors occur with equal probability at all sites, but the error containing proteins are then cleared from the cell with differing rates. We therefore analysed evolutionary substitution patterns, focusing on tyrosine codons as a randomly chosen example. We detected three separate substitutions for tyrosines on codon 172 (Y—F, Y—M and Y—W), as well as one (Y→C) on codon 206. In contrast, we do not observe any substitutions for the remaining four tyrosines contained within the detected peptides. An analysis of 50 homologous GST sequences reveals that position 172 shows much higher evolutionary variability than the other tyrosine positions, followed by position 206 (figure 3C). Moreover, although position 206 is much more highly conserved than 172, cysteines are tolerated in this position in evolution. These findings indicate that detectable substitutions are filtered by the effect they have on a protein’s structure, and we therefore detect substitutions predominantly in such positions and of such types that are structurally well tolerated.

Overall, these analyses lead us to conclude that error peptides are detected in nonrandom patterns, and these non-random patterns can be explained by the expected biological properties of such peptides.

### Quantitative patterns of substitutions depend on expression host and sequence

Although our mass spectrometric data do not allow comparisons of abundance between different peptides, relative changes in the spectral signal for the same peptide in different samples should be an accurate guide to changes in the frequency of the corresponding substitutions. The summed frequency of all substitutions showed strong differences for protein expressed from the two *E. coli* alleles (figure 4A), with protein expressed from the *disec* allele showing a higher cumulative signal than protein expressed from the *optec* allele. This is consistent with the view that non-optimal codons should show higher error rates than optimal ones. We observed this pattern for the P1, P2 and P3 peptides highlighted in figure 2 as well as the N-terminal peptide, and in the overall levels of substituted peptides for the entire protein. For reasons we cannot explain, the C-terminal P4 peptide showed substitution levels where this pattern appeared inverted.

**Figure 4.**
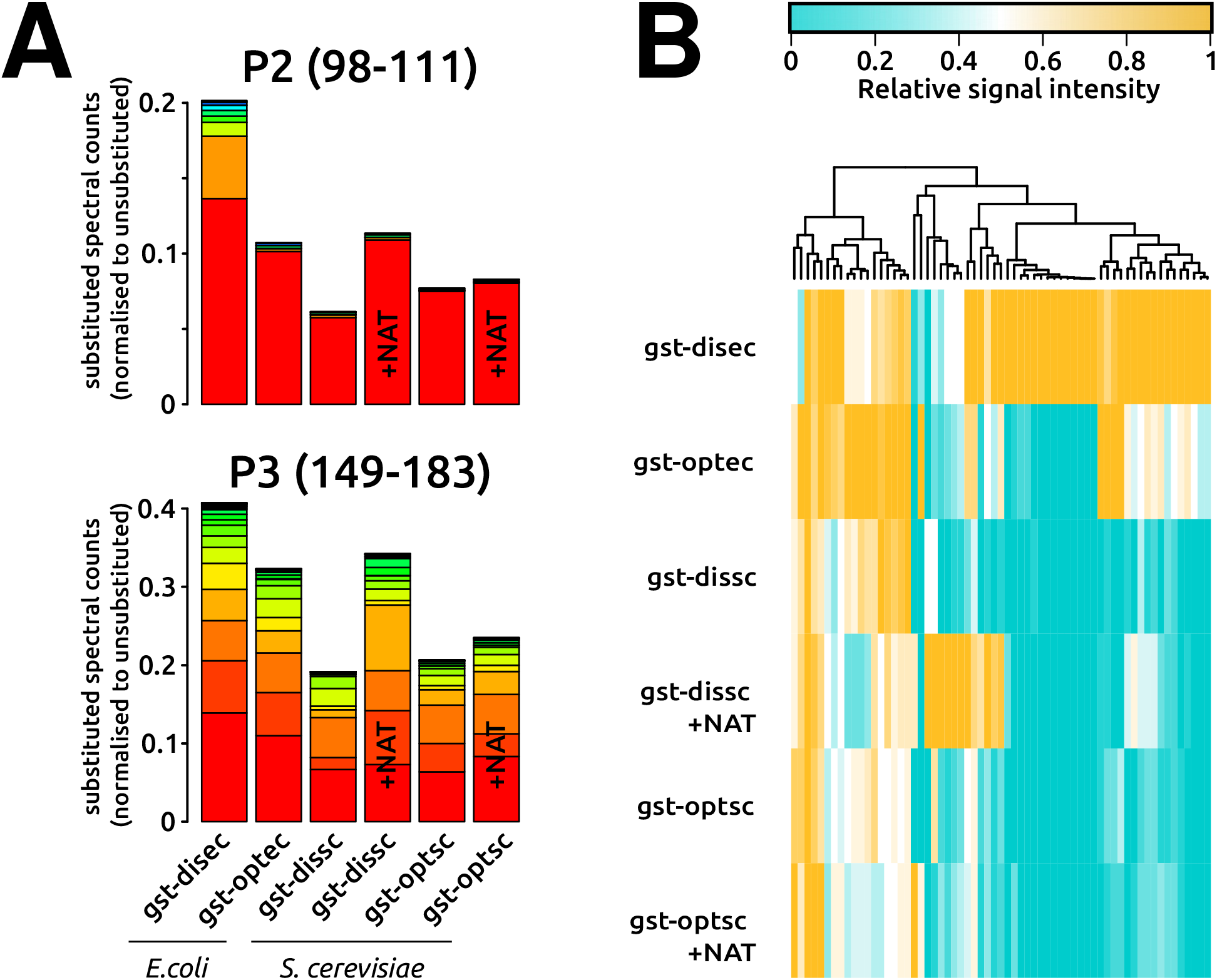
Quantitative changes in observed amino acid substitutions as a function of coding sequence. **A**, spectral counts (relative to counts for the corresponding wild-type peptides) for substitutions observed in samples generated from different coding sequences and in different expression hosts. +NAT, expression in the presence of 2 μg/ml nourseothricin. Within each graph the same colour indicates the same error peptide. **B**, relative detection of peptides in the different samples.

While the levels of error peptides differed strongly between the two *E. coli*-expressed genes, we observed less pronounced differences between the two yeast-expressed genes, and for the two peptides shown in figure 4A, the optimal construct actually generated higher error peptide signals than the dis-optimised one. To test whether this reflected genuine error levels or an inability to detect errors accurately in yeast-expressed proteins, we expressed the GST constructs in the presence of the error-inducing aminoglycoside, nourseothricin (NAT). We chose a NAT concentration of 2 μg/ml, where yeast growth is inhibited by 10-20% (data not shown). Especially for the *dissc* allele, samples expressed in the presence of the drug generate substantially higher error peptide signals than samples generated without the drug (figure 4A). Thus, our approach does detect substitutions in yeast proteins quantitatively, and the different allele specificity of errors in yeast and *E. coli* is likely due to genuine organismal differences.

Because the cumulative signal analysed in figure 4A is strongly influenced by peptides that give disproportionally high spectral signals, we also analysed the relative distribution of peptides between the different samples (figure 4B). Many of the individual error peptides generate the highest signal in the bacterial proteins, and most of these are preferentially detected with the *disec* allele. Such allele dependence is expected for codon misreading errors, but not for mis-acylation or transcriptional errors. Preferential misreading of the *disec* allele could be explained either if the misreading tRNA showed more stable base pairing patterns, or if the quantitative ratio between the misreading and cognate tRNAs was higher for this allele. We therefore systematically explored the base-pairing potential for the closest matching misreading tRNAs with the codons used in the two alleles (figure 5), for all substitutions where the *disec* allele generated at least a two-fold higher signal than the *optec* allele, and where this signal lay above a minimum threshold of 3000 units (this cut-off was intended to avoid problems with poor quantitation in the low signal range). Of the 13 substitutions in this category, 12 conformed to the expectation that the substituting tRNA can establish base-pairing patterns with the *disec* allele, which is a minimum condition for codon misreading to occur (figure 5). For nine of these 12 substitutions, either the base-pairing pattern was predicted to be weaker for the *optec* allele, or the tRNA ratio favoured substitutions for the *disec* allele (or both), thus potentially explaining the preferential misreading of this allele. Of the remaining four substitutions, one showed no predicted difference between the two alleles, and three showed predicted preferential interactions with the *optec* allele. In addition, for the Pro→Glu substitution no base pairing is predicted between the bacterial glutamyl tRNA anticodon and the proline codon used in the *disec* allele, so that this substitution is unlikely to be a genuine misreading error.

**Figure 5.**
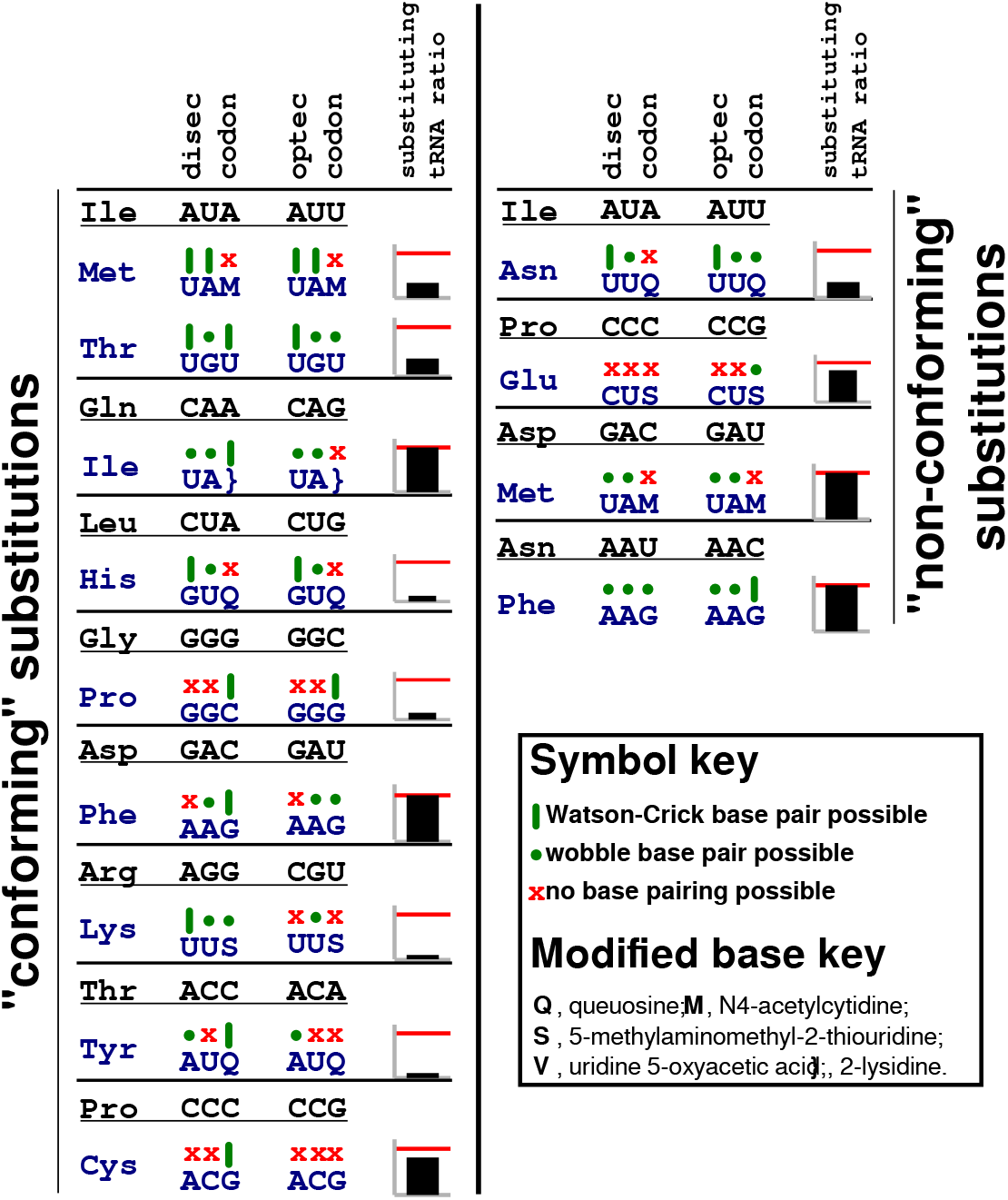
tRNA base-pairing patterns partially explain sequence-dependent amino acid substitution in bacteria. The base-pairing potential is shown for codons and the closest matching tRNA anticodon for substituted amino acids, for all substitutions which showed at least 2-fold higher abundance in the *gst-disec* allele compared to the *gst-optec* allele (note that anticodons are shown in 3’-5’ direction). Such allele-specific preferences could Į be explained if substitution are caused at the level of codon misreading, and if the misreading tRNA either pairs more stably with the *disec*-specific codon, or if its abundance ratio with the cognate tRNA is higher (or both). “Substituting tRNA ratio” displays the misreading tRNA:cognate tRNA ratio of the *gst-optec* codon relative to the *gst-disec* codon, the red line indicates a ratio of one (ie both alleles have the same ratio). tRNA abundances for these analyses are from ref^46^. “Conforming substitutions” show the expected differences in base-pairing potential and/ or tRNA ratios, whereas “non-conforming substitutions” do not. Base modifications are abbreviated as in the Modomics database^47^.

We observed five substitutions on codons for Met and Trp, which are encoded by a single codon and where the two alleles are therefore identical. For four of these substitutions, signals for the two alleles were within less than 30% of each other, and only one substitution (Met→Tyr) showed strong allele differences. Overall, these analyses show that many but not all of the allele-specific substitutions in the bacterially expressed proteins can be understood in terms of known features of the *E. coli* tRNA pool. The origin of the minority of substitutions which cannot be understood in these terms remains to be elucidated.

While none of the error peptides derived from yeast-expressed proteins showed clear allele differences, the signal for a number of peptides was altered in the presence of nourseothricin (NAT). To explore whether this dependence could also be explained by known properties of the tRNA pool, we divided the error peptides that were detected with a minimum threshold of 3000 units into those that showed at least a 2-fold higher signal in the presence of the drug (“NAT stimulated”), and those where the signal in the presence and absence of the drug differed by less than 30% (“non stimulated”). We then analysed how codons used in the *dissc* allele, which showed the highest degree of nourseothricin stimulation, base-paired with the anticodons of potentially substituting tRNAs (figure 6). Interestingly, all tRNA:codon combinations in the NAT stimulated pool were able to form extensive codon:anticodon base-pairs, with a high prevalence of non-Watson:Crick base pairs. In contrast, the anticodon:codon pairings in the “nonstimulated” set preferentially showed Watson:Crick or non-pairing base combinations. The mode of action by which nourseothricin induces misreading in eukaryotes is not understood in detail, but these results indicate that it may specifically increase the tolerance of the ribosomal A-site for non-Watson Crick contacts.

**Figure 6.**
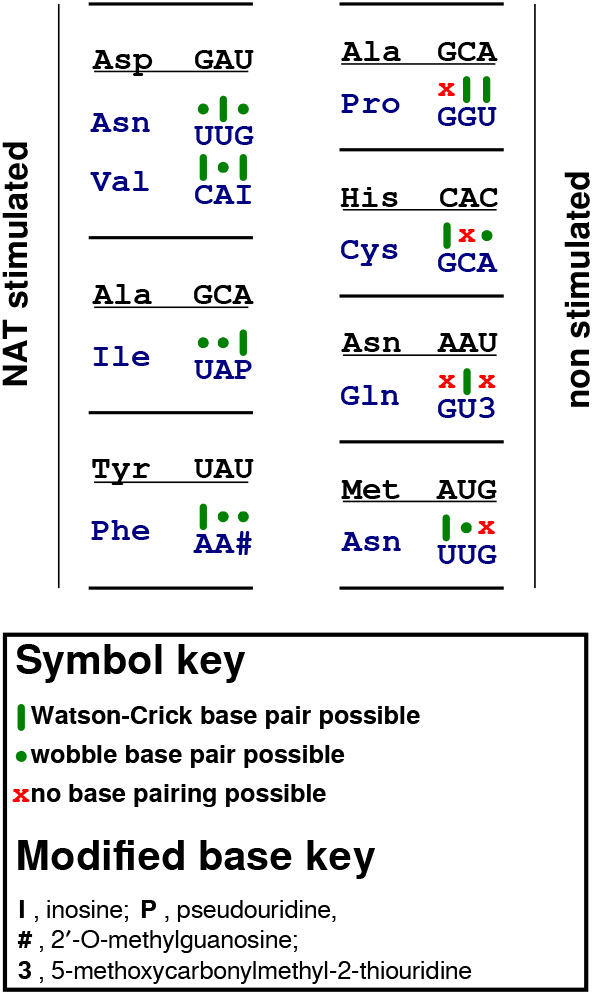
Stimulation of amino acid misincorporation by nourseothricin. Base pairing patterns are analysed for yeast codons used in the *dissc* allele, and the closest matching tRNA corresponding to observed substituted amino acids. The “NAT stimulated” substitutions (left) occur with signals that are at least twofold higher in the NAT-treated sample, whereas for the “non stimulated” substitutions (right) signals with or without NAT are within 30% of each other. tRNAs involved in NAT stimulated substitutions display more extensive base pairing patterns involving higher levels of non-Watson:Crick base pairs. Base modifications are abbreviated as in the Modomics database^47^.

### Correlation between error peptide abundance and enzymatic activity

The results discussed above indicate that codon optimisation may be particularly important in *E. coli* to avoid the generation of aberrant translation products, but less or not at all so in yeast. In order to further corroborate these findings, we investigated whether the different levels of error peptides correlated with differences in specific activity of the expressed proteins. Both the ability to bind immobilised glutathione and the ability to enzymatically catalyse transfer of glutathione onto a chromogenic substrate differed substantially between proteins expressed from the two *E. coli* alleles, but not between products of the two yeast alleles (Figures 7A and B). The activity differences thus inversely mirror the different levels of substitutions we observed, suggesting that a significant proportion of the detected substitutions may affect GST function.

**Figure 7.**
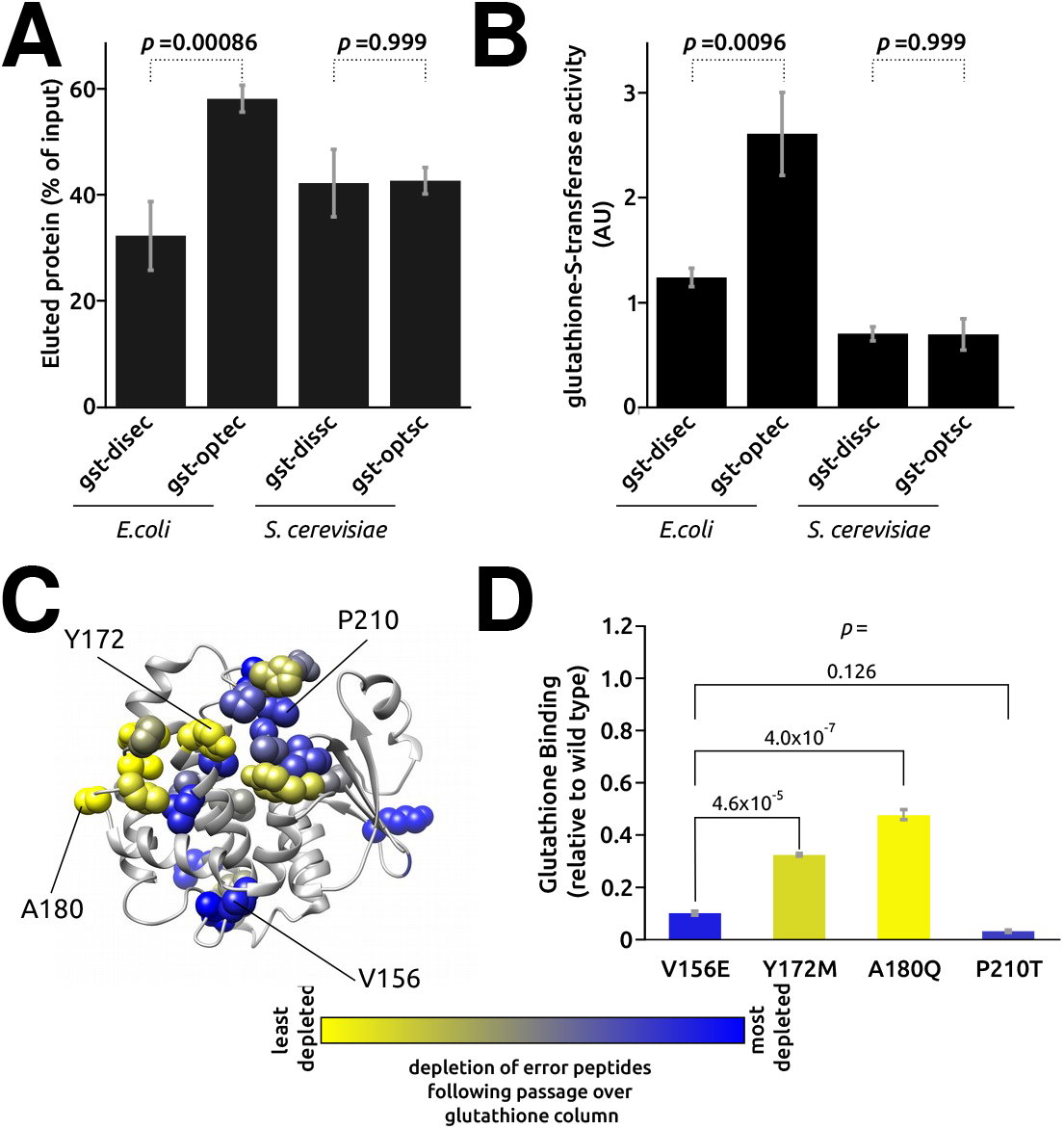
Differences in protein activity correlate with observed substitution levels. **A**, quantitative comparison of GST binding to immobilised gluathione. **B**, quantitative comparison of glutathione-S-transferase activity. Both assays show differences in specific activity of GST expressed from the bacterial alleles, which correlated inversely with the levels of substitutions observed in the proteins. **C**, depletion of specific substitutions in the gst-disec sample following passage over a glutathione column. **D**, mutant proteins mimicking four detected substitutions show differences in their ability to bind glutathione that correlate with the degree of depletion observed in panel C.

We directly tested this assumption by analysing samples of *disec* GST before and after passage over a glutathione affinity column. Levels of individual error peptides were strongly altered following passage over the column, with an up to 50-fold difference in spectral signal (figure 7C). There was a significant increase in the degree of depletion the closer the observed substitutions occurred to the glutathione binding site (p = 0.006, rho = 0.53 by Spearman’s rank correlation coefficient), likely reflecting a higher likelihood of substitutions near the binding site to affect the interaction between GST and glutathione. To test the veracity of this assay we selected two each of the most and least strongly depleted substitutions, and generated corresponding genetic mutants by site-directed mutagenesis of the bacterial expression vector. The glutathione binding activity of these mutants correlated strongly with the affinity inferred for the corresponding error products, and recovery of the two less depleted mutants differed from the two more depleted ones with high statistical significance (figure 7D). Thus, we have demonstrated experimentally that physiological translational errors in *E. coli* impact on protein function. The strong correlation between apparent amino acid misincorporation levels and protein activity also serves to corroborate the general validity of our mass spectrometric assays.

## Discussion

Our mass spectometry approach uncovered a surprising richness of detectable peptides with apparent amino acid substitutions, especially in proteins expressed in *E. coli* cells. Our initial analyses were designed to test how likely these events reflect genuine gene expression errors in the cell. By a number of criteria signals for these peptides behave as expected for such errors: their signal is always lower than signals of the corresponding error free peptides (figure 2B), they occur in patterns that are unlikely to arise by chance (figure 3A), the presence or absence of signals for specific modifications can be explained by examining the tolerance for corresponding mutations in evolution (figure 3C), and error peptide signals are increased by known error-inducing drugs (figure 4A). The analyses relating DNA sequence specific substitutions in bacteria to tRNA base pairing patterns (figure 5) further showed that many errors are potentially explicable in terms of molecular mechanisms, although for some this is not the case. Thus, while data for individual error peptides should for the time being be interpreted with caution, the sum of our data do allow forming strong conclusions on protein-wide error patterns. Previous studies have investigated tRNA-dependent codon misreading in *E. coli* in detail using luciferase-based reporter systems ^11,29,30^. Unfortunately we do not observe the substitutions analysed in depth in these studies in our GST reporter, possibly because they destabilise the GST protein and thus are depleted from the expressed pool. The only exception is a Phe181→Tyr recoding event which we detect with high signal in all analysed samples, and high Phe→Tyr recoding was also observed by Manickam *et al*. in their luciferase assays ^29^. Phe→Tyr recoding in the luciferase was not affected by ribosomal accuracy mutations, leading the study authors to conclude that this was unlikely to be a codon misreading event. Consistent with this we observe no sequence dependence of this event in *E. coli* and no responsiveness of the signal to nourseothricin. The uniformly high signal across all conditions and species indicates that this may be a mis-acylation event of phenylalanyl-tRNAs with physically similar tyrosine moieties. Thus, for the one substitution observed in both the luciferase-based work and in our study, the observed patterns are remarkably similar.

Based on the luciferase analyses the authors suggested that codon misreading errors depend on both the levels of competing tRNAs^11^ and on the strength of their interaction with the codon^30^. These principles are clearly reflected also in our findings. For those substitutions that are sequence dependent in *E. coli*, or that are stimulated by nourseothricin in yeast, codon misreading rather than misacylation or transcriptional errors is the most likely cause. For the majority of these substitutions, we observe substantial potential for base pairing between the codon and at least one tRNA delivering the substituted amino acid (figure 5 and 6). Moreover, for most of the sequence dependent substitutions in *E. coli*, we predict quantitative changes either in the strength of the base pairing or in the levels of the competing tRNAs which could explain the observed sequence dependence. Our work therefore corroborates the general principles uncovered by the luciferase studies, based on a much broader sample of tRNA:codon combinations.

It is interesting to compare the two biological systems used for expression of our model protein. Direct comparisons of luciferase-based assays for codon misreading by tRNA^Lys^ uuu have previously shown lower fidelity in *E. coli* than in yeast^20^, despite the fact that specialised ribosomal proofreading mechanisms able to remove translational error products were observed in *E. coli*^31^ but not in yeast^32^. Our data confirm the luciferase-based observations for amino acid misincorporation errors more generally (figure 4A). Moreover, the frequent detection of sequence specific misincorporation errors in *E. coli* is in stark contrast with the almost total absence of such sequence dependence in yeast (figure 4B), and we observe correlated effects on protein activity in *E. coli* but not in yeast (figure 5A,B). We explored whether more active surveillance mechanisms in yeast could potentially explain these patterns, but we failed to detect any evidence of turnover of significant levels of the yeast-expressed GST proteins in mutants of any of the known surveillance pathways and protein degradation mechanisms (data not shown). Shah and Gilchrist^25^ observed that the general occurrence of sequence-dependent errors depended on the exact balance of different tRNAs in the decoding population, and in particular on whether levels of cognate and near cognate tRNAs are correlated. It is not possible to rigorously evaluate *E. coli* or yeast tRNA populations in these terms because we do not understand well what constitutes near cognate tRNAs, but our data suggest that this correlation may be higher in yeast than in *E. coli*.

Lastly, our study demonstrates for the first time experimentally that bulk protein quality can be measurably affected by translational errors. Codon-dependent changes in protein quality have been observed in other studies^33^ as a function of ribosome speed and the ability of the nascent protein to fold, but we can rule out such effects in our case because the recombinant protein is more active when expressed from the more rapidly decoded *optec* allele (figure 6A,B). Instead, activity in our case correlates with and is likely directly affected by amino acid misincorporation errors. Evolutionary studies have long inferred that translational errors affect protein function sufficiently strongly to generate selective pressure against the use of error-prone codons ^14^. Our data support this albeit in a manner that depends on which organism is being considered.

## Materials and Methods

### DNA *constructs*

– Plasmids used in this study are listed in table 1. All recombinant DNA constructs were generated by ligation of restriction-enzyme treated DNA fragments *in vitro* and transformation into *E. coli* K12 JM109 ^34^.

**Table 1.**
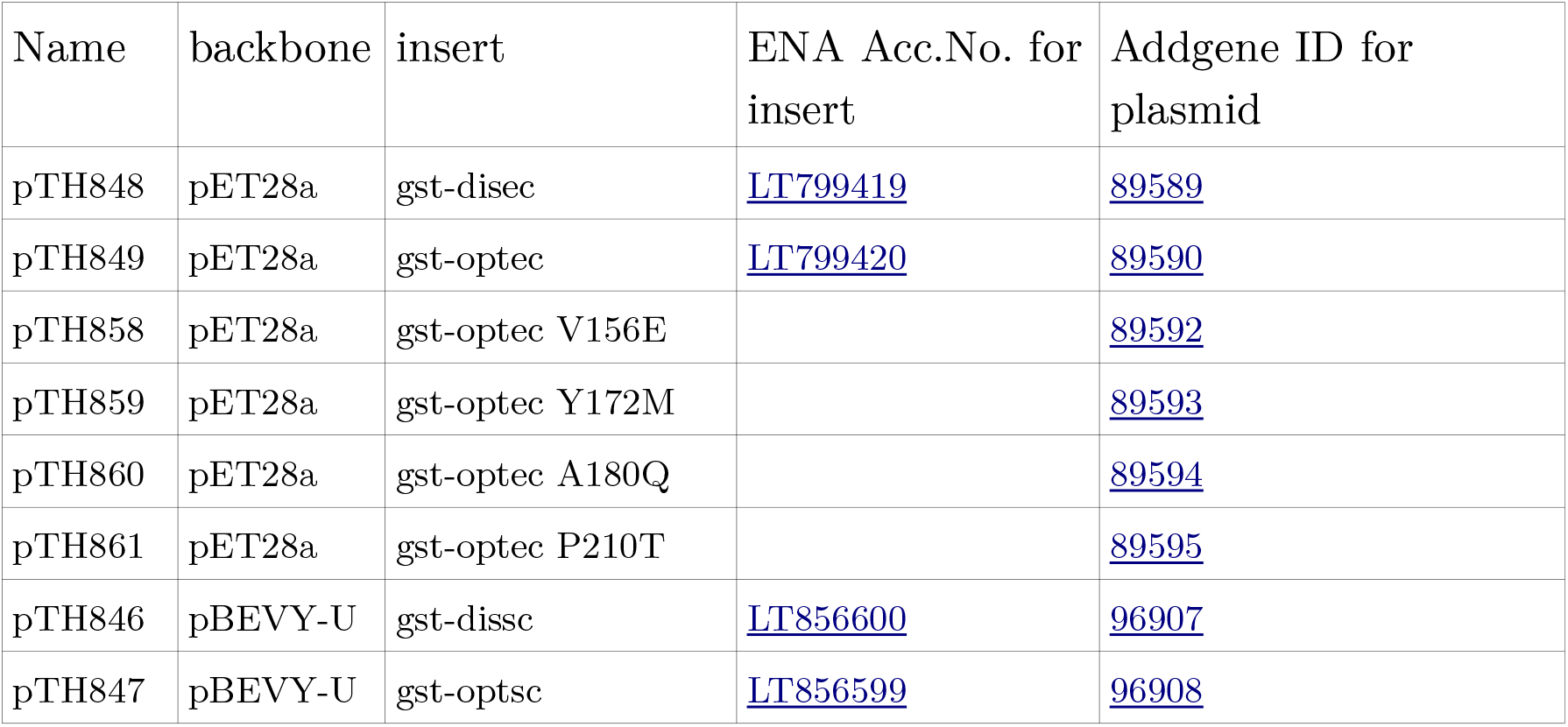
Plasmids used in this study.

DNA sequences were designed based on the GST protein sequence as used in the pGEX-series of expression vectors ^35^, preceded by eight histidine codons and an ATG start codon for the yeast sequences, or ATG and GGC codons (to enable cloning using the *NcoI* site in pET28a) for the bacterial constructs. For expression in *E. coli*, two coding sequences were designed by using exclusively the least or most preferred codons according to published *E. coli* codon usage tables ^23^. The complete synthetic sequences have been deposited in the European Nucleotide Archive^ii^ under accession numbers <u>LT799419</u> (gst-disec) and <u>LT799420</u> (gst-optec). For expression in *S. cerevisiae*, two coding sequences were designed consisting exclusively of the fastest or slowest decoded codons in this organisms, based on our most recent published models of codon decoding ^24^. The complete synthetic sequences have been deposited in the ENA under accession numbers <u>LT856600</u> (gst-dissc) and <u>LT856599</u> (gst-optsc).

DNA constructs were synthesized by Genscript (Piscataway, New Jersey, USA). For generation of the bacterial expression constructs, the synthetic genes were excised as *NcoI/HindIII* fragments from the supplier vector and ligated into *NcoI/HindIII* digested pET28a vector (Merck Millipore, UK). Point mutants in the GST coding sequence were constructed in the same vector using a PCR-based procedure ^36^. For generation of the yeast expression vectors, the synthetic genes were amplified by PCR using primers that introduced a *BamHI* site upstream of the start codon and an *XbaI* site downstream of the stop codon, and also removed a C-terminal arginine tag that had been present in the original synthetic construct. The PCR products were then cloned into *BamHI/XbaI* digested pBEVY-U ^37^.

### Bacterial protein expression

– Proteins were expressed in *E. coli* B BL21(DE3)^38^ transformed with the pET28a-based vectors. Cells were grown at 30°C in LB medium (2% Trypton, 1% Yeast Extract, 2% NaCl)) in the presence of 20 mg/l kanamycin to an oD_600_ of 0.8, induced by addition of IPTG to 1 mM final concentration, and incubated for a further 3 hours. Cells were pelleted by centrifugation at 15,000 rpm for 15 minutes and either frozen or used immediately for purification of the proteins.

Cell extracts were prepared by resuspending pellets in 5 ml/ g wet cell weight of lysis buffer (50mM Tris-HCl pH 7.5, 250 mM NaCl, 8 M Urea, 10 mM imidazole, 1x Protease Inhibitor Complete Mini EDTA-free [Sigma-Aldrich, UK]), and cells lysed by sonication. The cell extract was cleared by centrifugation for 15 minutes at 18,000 g before application to an IMAC column.

### Yeast protein expression

– Proteins were expressed in *S. cerevisiae* BY4741^39^ transformed with the pBEVY-U based vectors. Cells were grown in SC-Ura (0.67% yeast nitrogen base w/out amino acids, 2% glucose, 1.9 g/l Kaiser dropout mixture without uracil [Formedium, UK]), by inoculating from an overnight culture to a starting oD of 0.1, and growth at 30°C to a final oD of 2.0. Cells were pelleted by centrifugation at 15,000 rpm for 15 minutes, resuspended in 5 ml per gram wet cell weight Y-PER lysis buffer (Thermo Fisher Scientific, UK) which had been supplemented with Urea to 6M to create denaturing conditions, agitated at room temperature for 30 minutes, and the lysate then cleared by centrifugation at 19,000 g for 20 minutes.

### Protein purification

– Cell lysates from cells grown in 0.5 to 2 l of culture were passed over 1 ml of Chelating Sepharose Fast Flow (GE Healthcare, UK) which had previously been charged with NiCl_2_ and equilibrated with lysis buffer. The resin was then washed with 10 ml renaturing buffer (50 mM Tris-HCl pH 7.5, 100 mM NaCl, 5 mM 2-mercaptoethanol) and protein eluted with elution buffer (renaturing buffer also containing 100 mM EDTA).

### Western blotting

– SDS-PAGE and western blotting were performed as described^40^. Anti-GST antibody was from Sigma-Aldrich (G7781, Sigma-Aldrich, UK) and was used at a dilution of 1:1000, with HRP-labelled anti-rabbit IgG (12-348, Sigma-Aldrich, UK, 1:15000) as secondary antibody.

### Glutathione binding assays

– Purified GST was brought to a concentration of 0.2 mg/ml using 10,000 MWCO Centrifugal Concentrators. Glutathione Magnetic Agarose Beads from 100 μl of slurry (Pierce, UK) were washed with 2 x 1ml Buffer (50 mM Tris-Hcl pH 7.8, 100 mM NaCl, 5 mM MgCl_2_, 5 mM 2-mercaptoethanol) before adding 0.5 ml of the adjusted GST samples and agitating for 15 minutes at room temperature. Supernatant was removed and retained as the “unbound” sample. Beads were washed twice with 1ml buffer, bound protein eluted in 200 μl SDS-PAGE sample buffer at 95°C for 10 minutes, and the unbound and eluted protein fractions analysed by SDS-PAGE and western blotting with anti-GST antibodies.

### Glutathione-S-transferase assay

– A commercial Glutathione-S-Transferase Assay Kit (CS0410, Sigma-Aldrich, UK) was used to determine the specific enzymatic activity of purified GST samples in 96-well format. 2 μl of protein diluted to approximately 0.1 μg/μl in elution buffer, 18 μl of the sample buffer provided with the kit, and 180 μl of master mix were mixed in the wells of a clear 96-well plate. Absorbance at 340 nm was measured every 30 seconds for 20 minutes. Absorbance was plotted against time, and slopes of the linear parts of the plots determined. For normalisation against actual protein content, samples were removed from the wells and transferred onto PVDF membrane using a slot blot device. The relative amount of membrane-bound GST was determined using anti-GST antibody as described above for western blotting.

### Tryptic digest

– The procedure was adapted from a published protocol ^41^. The preparation of the gel bands involved dicing, washing, removal of Coomassie Blue, reduction and alkylation of cysteines, digesting overnight at room temperature in a solution containing 10 ng/ μl trypsin, and sonication of the gel fragments to release the tryptic peptides.

### Mass spectrometry

– Peptide mixtures were analysed by ultra performance nanoLC (ACQUITY M Class, Waters) coupled to an IMS mass spectrometer (SYNAPT G2-Si, Waters) fitted with a NanoLockSpray source (Waters). Samples were loaded via a Symmetry C18 5 μm, 180 μm x 20 mm trap column (Waters) and separated through a HSS T3 C18 1.8 μm, 75 μm x 150 mm analytical column (Waters). Peptides were eluted using a 3 % to 40 % acetonitrile 0.1 % formic acid gradient over 40 min at a flow rate of 300 nL/min. The mass spectrometer was operated in positive ion mode with a capillary voltage of 3.25 kV, cone voltage of 30 V and a source offset of 80 V. Before analysis the instrument was calibrated with NaI and during analysis a LockMass reference, Glu-1-fibrinopeptide B (Waters), was delivered to the NanoLockSpray source. Mass spectra were collected, over 50–2000 m/z, alternating between low (4 eV) and elevated (15–45 eV) collision energies at a scan speed of 0.5 s.

High Definition MSe (HDMSe) spectra were LockMass corrected and aligned using Progenesis QIP (Nonlinear Dynamics) before further processing to deconvolute parent-fragment peak lists. Peak lists were searched against a custom database, which was generated as described in the main text. The search was conducted with the following ion-matching requirements: matching intact mass and at least 3 fragments per peptide, 5 fragments per protein and 1 peptide per protein (the latter two settings are meaningless given that the peptide database consisted of individual tryptic peptides, but had to be set in the search software). Enzyme specificity was set to trypsin allowing for 1 missed cleavage, and with modifications of carbamidomethyl of cysteine (fixed) and oxidation of methionine (variable). Only data for peptides detected with a score > 5 were used in subsequent analyses.

*Computational analyses* – we used the ConSurf server^42^, with automatic sampling of 50 GST homologues from the UNIREF-90 database and otherwise default settings, for evaluating evolutionary conservation of residues (figure 3). To analyse the degree of correlation between peptide depletion and distance of the substituted amino acid from the glutathione binding site (figure 7), distances between the C**α** atom of Tyr15 (one of the residues in direct contact with glutatione ^43^) and the C**α** atoms of the substituted residues were determined using UCSF Chimera ^44^. All other analyses were conducted using R^45^. Where statistical significance is indicated, tests were based on ANOVA and post-hoc analyses using Tukey’s Honestly Significant Difference criteria.

## Acknowledgements

This study was funded by a Proof of Concept grant awarded to TVDH through BioProNET, a BBSRC NIBB co-sponsored by the EPSRC (BBSRC ref BB/L013770/1).

## Footnotes

i Due to the cloning procedures used, the *E. coli* sequences contain a single additional glycine between the N-terminal methionine and the His-tag. Where amino acid numbers are given in the manuscript, these refer to the yeast sequences (or bacterial sequences - 1).

ii www.ebi.ac.uk/ena/

## References

1. Ribas de Pouplana, L., Santos, M. A. S., Zhu, J.-H., Farabaugh, P. J. & Javid, B. Protein mistranslation: friend or foe? Trends Biochem. Sci. 39, 355–362 (2014).

2. Gingold, H. & Pilpel, Y. Determinants of translation efficiency and accuracy. Mol. Syst. Biol. 7, 481 (2011).

3. Guo, H. H., Choe, J. & Loeb, L. A. Protein tolerance to random amino acid change. Proc. Natl. Acad. Sci. U. S. A. 101, 9205–9210 (2004).

4. Ruan, B. et al. Quality control despite mistranslation caused by an ambiguous genetic code. Proc. Natl. Acad. Sci. U. S. A. 105, 16502–16507 (2008).

5. Kalapis, D. et al. Evolution of Robustness to Protein Mistranslation by Accelerated Protein Turnover. PLOS Biol. 13, e1002291 (2015).

6. Reynolds, N. M. et al. Cell-specific differences in the requirements for translation quality control. Proc. Natl. Acad. Sci. U. S. A. 107, 4063–4068 (2010).

7. von der Haar, T. et al. The control of translational accuracy is a determinant of healthy ageing in yeast. Open Biol. 7, 160291 (2017).

8. Liu, Y. et al. Deficiencies in tRNA synthetase editing activity cause cardioproteinopathy. Proc. Natl. Acad. Sci. 111, 17570–17575 (2014).

9. Vermulst, M. et al. Transcription errors induce proteotoxic stress and shorten cellular lifespan. Nat. Commun. 6, 8065 (2015).

10. Ling, J., Reynolds, N. M. & Ibba, M. Aminoacyl-tRNA Synthesis and Translational Quality Control. Annu. Rev. Microbiol. 63, 61–78 (2009).

11. Kramer, E. B. & Farabaugh, P. J. The frequency of translational misreading errors in E. coli is largely determined by tRNA competition. RNA 13, 87–96 (2007).

12. Wohlgemuth, I., Pohl, C. & Rodnina, M. V. Optimization of speed and accuracy of decoding in translation. EMBO J. 29, 3701–3709 (2010).

13. Ke, Z. et al. Translation fidelity coevolves with longevity. Aging Cell 1–6 (2017). doi:10.1111/acel.12628

14. Zhou, T., Weems, M. & Wilke, C. O. Translationally optimal codons associate with structurally sensitive sites in proteins. Mol. Biol. Evol. 26, 1571–80 (2009).

15. de Nadal, E., Fadden, R. P., Ruiz, a, Haystead, T. & Ariño, J. A role for the Ppz Ser/Thr protein phosphatases in the regulation of translation elongation factor 1Balpha. J. Biol. Chem. 276, 14829–34 (2001).

16. Röther, S. & Strässer, K. The RNA polymerase II CTD kinase Ctk1 functions in translation elongation. Genes Dev. 21, 1409–1421 (2007).

17. Stansfield, I. et al. Missense translation errors in Saccharomyces cerevisiae. J. Mol. Biol. 282, 13–24 (1998).

18. Stahl, G., Bidou, L., Rousset, J. P. & Cassan, M. Versatile vectors to study recoding: conservation of rules between yeast and mammalian cells. Nucleic Acids Res. 23, 1557–1560 (1995).

19. Salas-Marco, J. & Bedwell, D. M. Discrimination between defects in elongation fidelity and termination efficiency provides mechanistic insights into translational readthrough. J. Mol. Biol. 348, 801–815 (2005).

20. Kramer, E. B., Vallabhaneni, H., Mayer, L. M. & Farabaugh, P. J. A comprehensive analysis of translational missense errors in the yeast Saccharomyces cerevisiae. RNA 16, 1797–1808 (2010).

21. Mohler, K. et al. MS-READ: Quantitative Measurement of Amino Acid Incorporation. Biochim. Biophys. Acta - Gen. Subj. 1–8 (2017). doi:10.1016/j.bbagen.2017.01.025

22. Cvetesic, N. et al. Proteome-wide measurement of non-canonical bacterial mistranslation by quantitative mass spectrometry of protein modifications. Sci. Rep. 6, 28631 (2016).

23. Nakamura, Y., Gojobori, T. & Ikemura, T. Codon usage tabulated from international DNA sequence databases: status for the year 2000. Nucleic Acids Res. 28, 292 (2000).

24. Chu, D. et al. Translation elongation can control translation initiation on eukaryotic mRNAs. EMBO J. 33, 21–34 (2014).

25. Shah, P. & Gilchrist, M. A. Effect of correlated tRNA abundances on translation errors and evolution of codon usage bias. PLoS Genet. 6, (2010).

26. Burgess-Brown, N. A. et al. Codon optimization can improve expression of human genes in Escherichia coli: A multi-gene study. Protein Expr. Purif. 59, 94–102 (2008).

27. Bateman, R. H. et al. A novel precursor ion discovery method on a hybrid quadrupole orthogonal acceleration time-of-flight (Q-TOF) mass spectrometer for studying protein phosphorylation. J. Am. Soc. Mass Spectrom. 13, 792–803 (2002).

28. Silva, J. C. et al. Quantitative Proteomic Analysis by Accurate Mass Retention Time Pairs. Anal. Chem. 77, 2187–2200 (2005).

29. Manickam, N., Nag, N., Abbasi, A., Patel, K. & Farabaugh, P. J. Studies of translational misreading in vivo show that the ribosome very efficiently discriminates against most potential errors. RNA 20, 9–15 (2014).

30. Manickam, N., Joshi, K., Bhatt, M. J. & Farabaugh, P. J. Effects of tRNA modification on translational accuracy depend on intrinsic codon–anticodon strength. Nucleic Acids Res. gkv1506 (2015). doi:10.1093/nar/gkv1506

31. Zaher, H. S. & Green, R. Quality control by the ribosome following peptide bond formation. Nature 457, 161–6 (2009).

32. Eyler, D. E. & Green, R. Distinct response of yeast ribosomes to a miscoding event during translation. RNA 17, 925–32 (2011).

33. Hess, A.-K., Saffert, P., Liebeton, K. & Ignatova, Z. Optimization of Translation Profiles Enhances Protein Expression and Solubility. PLoS One 10, e0127039 (2015).

34. Yanisch-Perron, C., Vieira, J. & Messing, J. Improved M13 phage cloning vectors and host strains: nucleotide sequences of the M13mp18 and pUC19 vectors. Gene 33, 103–119 (1985).

35. Kaelin, W. G. et al. Expression cloning of a cDNA encoding a retinoblastoma-binding protein with E2F-like properties. Cell 70, 351–364 (1992).

36. Mikaelian, I. & Sergeant, A. A general and fast method to generate multiple site directed mutations. Nucleic Acids Res. 20, 376 (1992).

37. Miller, C. A. I., Martinat, M. A. & Hyman, L. E. Assessment of aryl hydrocarbon receptor complex interactions using pBEVY plasmids: expression vectors with bi-directional promoters for use in Saccharomyces cerevisiae. Nucleic Acids Res. 26, 3577–3583 (1998).

38. Studier, F. W. & Moffatt, B. A. Use of bacteriophage T7 RNA polymerase to direct selective high-level expression of cloned genes. J. Mol. Biol. 189, 113–130 (1986).

39. Brachmann, C. B. et al. Designer deletion strains derived from Saccharomyces cerevisiae S288C: a useful set of strains and plasmids for PCR-mediated gene disruption and other applications. Yeast 14, 115–132 (1998).

40. Antibodies: a laboratory manual. (Cold Spring Harbor Laboratory Press, 2014).

41. Shevchenko, A., Tomas, H., Havlis, J., Olsen, J. V & Mann, M. In-gel digestion for mass spectrometric characterization of proteins and proteomes. Nat. Protoc. 1, 2856–60 (2006).

42. Ashkenazy, H. et al. ConSurf 2016: an improved methodology to estimate and visualize evolutionary conservation in macromolecules. Nucleic Acids Res. 44, W344–50 (2016).

43. Cardoso, R. M. F., Daniels, D. S., Bruns, C. M. & Tainer, J. A. Characterization of the electrophile binding site and substrate binding mode of the 26-kDa glutathione S-transferase from Schistosoma japonicum. Proteins 51, 137–146 (2003).

44. Pettersen, E. F. et al. UCSF Chimera--a visualization system for exploratory research and analysis. J. Comput. Chem. 25, 1605–1612 (2004).

45. R Development Core Team. R: A Language and Environment for Statistical Computing. (R Foundation for Statistical Computing, 2015).

46. Dong, H., Nilsson, L. & Kurland, C. G. Co-variation of tRNA Abundance and Codon Usage in Escherichia coli at Different Growth Rates. J. Mol. Biol. 260, 649–663 (1996).

47. Czerwoniec, A. et al. MODOMICS: a database of RNA modification pathways. 2008 update. Nucleic Acids Res. 37, D118–D121 (2009).

